# SurfDock is a Surface-Informed Diffusion Generative Model for Reliable and Accurate Protein-ligand Complex Prediction

**DOI:** 10.1101/2023.12.13.571408

**Authors:** Duanhua Cao, Mingan Chen, Runze Zhang, Jie Yu, Xinyu Jiang, Zhehuan Fan, Wei Zhang, Mingyue Zheng

**Author notes:** **Corresponding Authors** (Mingyue Zheng). **Author Contributions** D.H.C., M.A.C., R.Z.Z. contributed equally. M.Y.Z. conceived the research project. D.H.C and M.A.C developed the method and implemented the code. All authors contributed to the analysis of the results. D.H.C., M.A.C., and M.Y.Z. wrote the paper. All authors read and approved the manuscript. **Notes** The authors declare no competing financial interest.

## Abstract

In the field of structure-based drug design, accurately predicting the binding conformation of ligands to proteins is a long-standing objective. Despite recent advances in deep learning yielding various methods for predicting protein-ligand complex structures, these AI-driven approaches frequently fall short of traditional docking methods in practice and often yield structures that lack physical and chemical plausibility. To overcome these limitations, we present SurfDock, an advanced geometric diffusion network, distinguished by its ability to integrate multiple protein representations including protein sequence, three-dimensional structural graphs, and surface-level details into its equivariant architecture. SurfDock employs a generative diffusion model on a non-Euclidean manifold, enabling precise optimization of molecular translations, rotations, and torsions for reliable binding poses generation. Complemented by a mixture density network for scoring using the same comprehensive representation, SurfDock achieves significantly improved docking success rates over all existing methods, excelling in both accuracy and adherence to physical constraints. Equipped with post-docking energy minimization as an optional feature, the plausibility of generated poses is further enhanced. Importantly, SurfDock demonstrates excellent generalizability to unseen proteins and extensibility to virtual screening tasks with state-of-the-art performance. We consider it a transformative contribution that could serve as an invaluable asset in structure-based drug design.

## INTRODUCTION

The realm of life sciences is currently undergoing a renaissance, sparked by groundbreaking advancements in artificial intelligence (AI), particularly deep learning (DL)^1-5^. This wave of technological innovation is redefining the landscape of structure-based drug design (SBDD), a pivotal domain in pharmaceutical research. SBDD hinges on the intricate understanding of protein-ligand interactions, with the objective to discover or design ligands that bind to specific proteins, thereby modulating their function for therapeutic purposes^6, 7^. Understanding these interactions is crucial because the effectiveness of drugs depend heavily on how well they bind to their target proteins, and whether they affect any other proteins in the body. Accurate and rapid prediction of protein-ligand complex structures is pivotal for virtual screening, a process that screens potential drugs from extensive databases against specific protein targets. To date, the advancement of high-throughput structure-based virtual screening techniques has significantly contributed to various drug discovery applications, notably accelerating the pace of drug discovery^8, 9^.

Nonetheless, predicting how a small molecule binds to a protein, often referred to as ligand docking, presents substantial complexity. This complexity arises from the dynamic and multifaceted nature of protein-ligand interactions. Ligand docking generally involves two phases: the generation of docking poses and their subsequent scoring^10^. The initial phase aims to identify feasible binding poses for ligands relative to target proteins, whereas the scoring phase involves evaluating these poses to estimate binding affinity. Traditional methods in ligand docking, such as AutoDock Vina^11^, Glide^12^, and Gold^13^ employ heuristic algorithms to explore potential ligand conformations. However, they often struggle to comprehensively cover the vast conformational space, potentially overlooking feasible binding poses. This incomplete exploration can result from their inherent algorithmic constraints, which prioritize computational speed over thoroughness^14^. The scoring algorithms in these traditional methods apply simplistic functional terms to estimate binding affinity of docked poses. Researchers have been working on improving scoring functions based on these traditional search techniques, like SMINA^15^, GNINA^16^, DeepDock^17^ and other machine learning scoring functions^18, 19^. While the subsequent scoring phase is also important, it relies on the quality of the generated poses^10, 20^. If the initial pose generation algorithm is flawed, even an accurate scoring system can be misled, leading to suboptimal ligand selections. This limitation is particularly evident in virtual screening contexts, where identifying suitable protein-ligand interactions and ligand conformations for a known protein’s binding pocket is crucial. As a result, developing efficient algorithms for ligand docking is of crucial importance.

This is where DL methods become particularly valuable. With the high-quality data available from sources like the Protein Data Bank (PDB), DL models can decipher the complex interplay between proteins and ligands^21^. This capability enhances the prediction accuracy of protein-ligand complex structures. For the pose generation task, pervious deep learning approaches like Uni-Mol^22^, EquiBind^23^, E3Bind^24^,TANKBind^25^ and KarmaDock^26^ predominantly treated it as a regression problem, predicting the binding pose of a ligand to a protein in a one-shot manner. Although these methods are faster, their accuracy improvements over classical methods have been limited. This limitation may stem from the inherent discord between the regression-centric approach and the actual process of ligands binding to the targets, i.e., the interactive process between ligands and the targets to find the most suitable binding conformations. In this context, works by Jaakkola et al. introduces a paradigm shift by treating molecular docking as a generative modeling problem^27^. Unlike regression methods, their work DiffDock learns a distribution over possible ligand poses. This approach is implemented through a diffusion generative model (DGM)^28^, which defines a diffusion process over the critical degrees of freedom in docking: translations, rotations, and torsions. In recent years, diffusion networks have demonstrated remarkable success in a variety of tasks related to molecular generation and conformer generation^29-31^. DiffDock’s methodology, emphasizing iterative refinement of ligand poses through updates in translations, rotations, and torsion angles from a noisy prior to a learned distribution, mirrors the complex nature of molecular interactions.

Despite these advancements, challenges persist. Studies by Ke et al. indicate that in practical SBDD tasks, where the binding pocket is known, DL methods have not yet outperformed traditional approaches^32^. Additionally, many AI-generated poses, though technically successful in terms of Root Mean Square Deviation (RMSD) metrics (i.e., if the RMSD between a generated ligand pose and a crystal ligand pose is less than 2Å, we consider the docking is successful^33^), exhibit biophysical inconsistencies, such as intermolecular steric clashes or unrealistic bonds or angles^34^. Thus, the over-reliance on RMSD for pose evaluation is increasingly acknowledged as inadequate, failing to capture the subtleties of molecular interactions and physical realities of binding poses. Recognizing these limitations, recent efforts have focused on developing better metrics for assessing the rationality of generated poses^34, 35^. Deane et al. introduced PoseBusters, a tool designed to evaluate poses based on physical and chemical rationality, and PoseBusters Benchmark set, a novel set of 428 complexes released from 2021 onwards^34^. Their findings suggest that, when considering the plausibility of generated poses, DL methods have not outperformed traditional techniques. Moreover, it shows that all DL methods perform poorly on proteins with less than 30% sequence similarity to the training set. These two findings suggest that it is challenging for current DL algorithms to generate biophysically plausible complex structures and to generalize to novel proteins. One possible reason for the current shortcomings of DL methods is their reliance on coarse-grained, residue-level representations of proteins. Ideally, a more accurate all-atom representation of the protein or its binding pocket would offer greater precision, but with substantial computational demands. The conventional coarse-grained representation tends to oversimplify protein structures, consequently expanding the ligand pose search space into regions already occupied by protein atoms, potentially resulting in intermolecular clashes. Recent studies, however, have demonstrated the benefits of incorporating protein surface-level information, which offers a more detailed representation by modeling proteins as continuous shapes with geometric and chemical properties^17, 36-41^. By utilizing this surface information to more accurate describe geometric space in binding pocket, we anticipate a reduction in the occurrence of intermolecular clashes. Additionally, successes in sequence-based drug design and protein structure prediction have highlighted the value of sequence information in protein representation^1, 3, 42-44^. Building on these insights, we hypothesized that by leveraging multimodal protein information and advanced generative modeling frameworks, it might be possible to address the current issues in molecular docking while maintaining computational efficiency.

In this work, we introduce SurfDock, a geometric diffusion network designed for generating reliable binding ligand poses. SurfDock is conditioned on the protein pocket and a random starting ligand conformation, and it includes an internal scoring module SurfScore trained on crystal protein-ligand complexes for confidence estimation. By incorporating multimodal protein information—surface features, residue structure features, and pre-trained sequence-level features—into a surface node level representation, SurfDock achieves top performance in docking success rates across several benchmarks, including PDBbind2020^45^, the Astex Diverse Set^46^, and the PoseBusters benchmark set^34^. When evaluating the plausibility of generated poses using the PoseBuster tool, SurfDock demonstrates a significant improvement in pose rationality compared to previous DL methods. Additionally, SurfDock incorporates an optional fast force field relaxation step for protein-fixed ligand optimization, further enhancing its accuracy. This improvement allows SurfDock to surpass all existing DL and traditional methods in both docking success rates and pose plausibility. Besides, we also find that SurfDock generalizes effectively to new proteins. In the latter part of our study, we conducted a comprehensive evaluation of SurfDock on the virtual screening benchmark dataset DEKOIS2.0^47^. Our results clearly demonstrate that SurfDock not only meets but exceeds the performance of existing docking methods in this domain. This performance, combined with its practicality and reliability, positions SurfDock as a valuable contribution to the SBDD community.

## RESULTS AND DISCUSSION

### Method Overview

Our ligand docking framework, SurfDock, comprises primarily two stages: a diffusion network for pose generation and a scoring module (SurfScore), supplemented by an optional post-docking energy minimization module. Both the generation and scoring modules employ identical protein-ligand representation layers.

For protein binding pocket representation, SurfDock utilizes a tri-level approach: sequence level, residue graph level, and surface level. For the first two levels, SurfDock employs residue structural features and embeddings from the large language model ESM-2^48^ for residue representation. Unique to SurfDock is the integration of a molecular surface representation of the binding site, formatted as a polygon mesh using MaSIF^37^. This mesh comprises nodes, edges, and faces that collectively define the molecular surface’s shape, with nodes encapsulating chemical and topological features and edges representing node connectivity. The sequence and residue graph embeddings are then mapped onto this molecular surface, as illustrated in **Figure 1 a**. Ligands in SurfDock are represented as 3D atomic-level graphs, where nodes symbolize atoms and edges denote expanded interatomic distances.

**Fig. 1:**
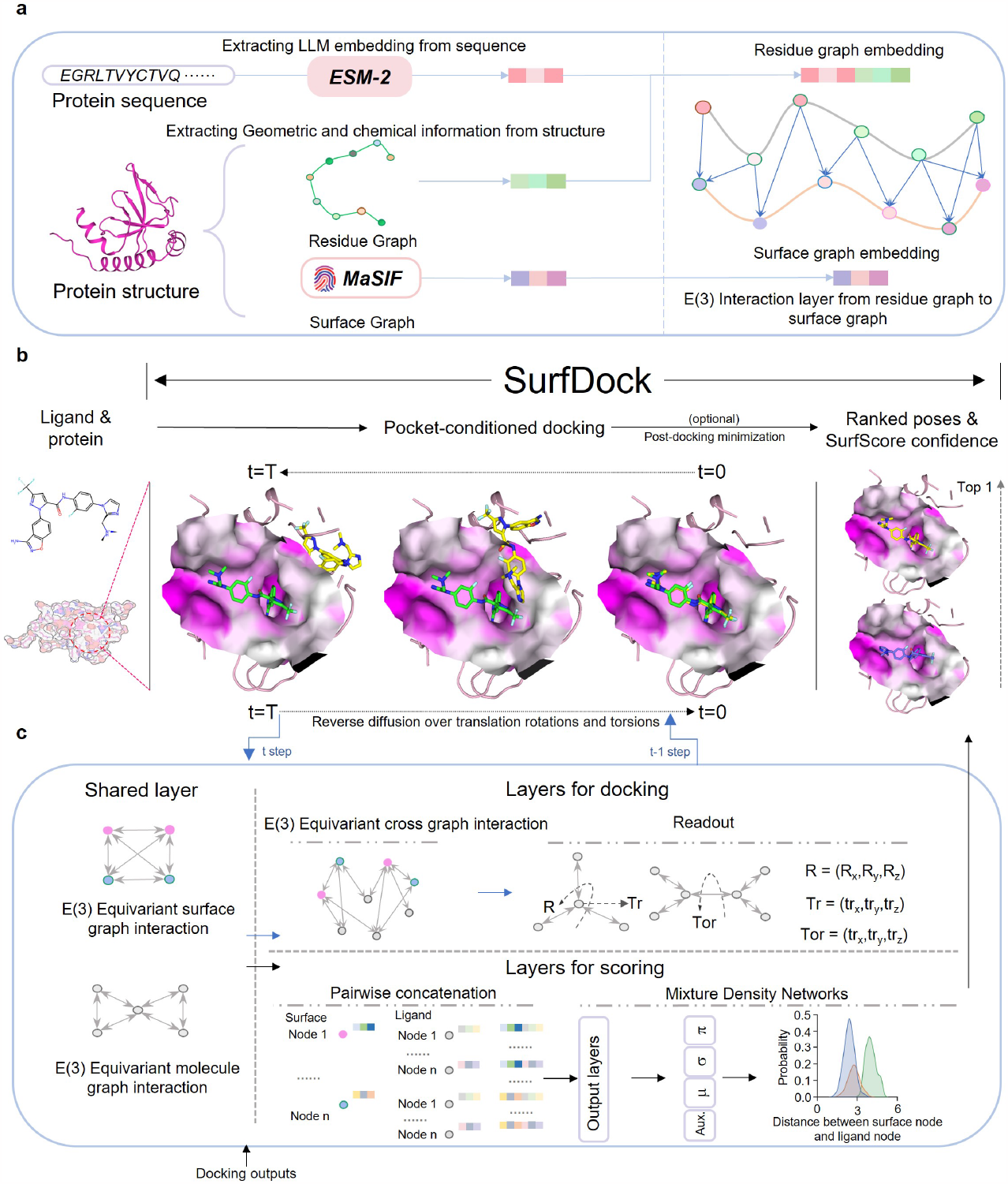
The overall architecture of SurfDock. **a**: Illustration of the multimodal representation of proteins in SurfDock. Embeddings from protein sequence and residue graph are mapped onto the surface graph. **b**: Overview of SurfDock. The model takes separate protein and ligand as inputs. Starting from a random initial ligand pose, SurfDock refines (or denoises) the pose over translational, rotational, and torsional degrees of freedom conditioned on the pocket. The output is complemented with a confidence score provided by SurfScore, with or without an optional energy minimization. **c**: The docking and scoring both uses the same representation of pocket and ligands, but different readout layers. This enables simultaneous pose generation and confidence estimation without additional scoring model.

Based on these representations, the geometric diffusion network in the first stage learns to refine (or denoise) a random initialized ligand pose conditioned on the binding pocket. To learn the distribution of protein-ligand complexes, we train the diffusion generation module using PDBbind2020 dataset, which contains experimental 3D data of ligands bound to protein targets and the binding affinities., with the protein’s binding pocket serving as a conditional factor for generating ligand poses. The diffusion process incrementally introduces noise into the ligand’s pose, encompassing translational, rotational, and torsional alterations while the generative process learns to reconstruct the ligand’s pose by refining a noise-altered structure back to its ground-truth conformation.

The scoring module SurfScore in the second stage aligns closely with our diffusion generation module in terms of representation. This integration marks a departure from previous deep learning approaches like DiffDock, which typically trained their pose generation and scoring modules separately with distinct training objectives. For instance, DiffDock’s scoring module was trained on a binary classification basis, using positive and negative samples produced by its pose generation module. Moreover, DiffDock used a coarse-grained representation for pose generation module and all-atom representation for scoring module. In contrast, SurfScore shares not only the representation layer with the generation module but also its training objective, focusing on the same crystal protein-ligand complexes and supplemented by a mixture density network^17, 49^ for scoring. This unified approach is designed to enhance the synergy between the pose generation and scoring stages, potentially leading to improved performance in ligand docking, as we aim to demonstrate in **Fig. 3**. Moreover, by utilizing a common representation and input for both modules, our method eliminates the need for separate pose generation, format conversion, and scoring processes, streamlining the entire pipeline.

**Fig. 2:**
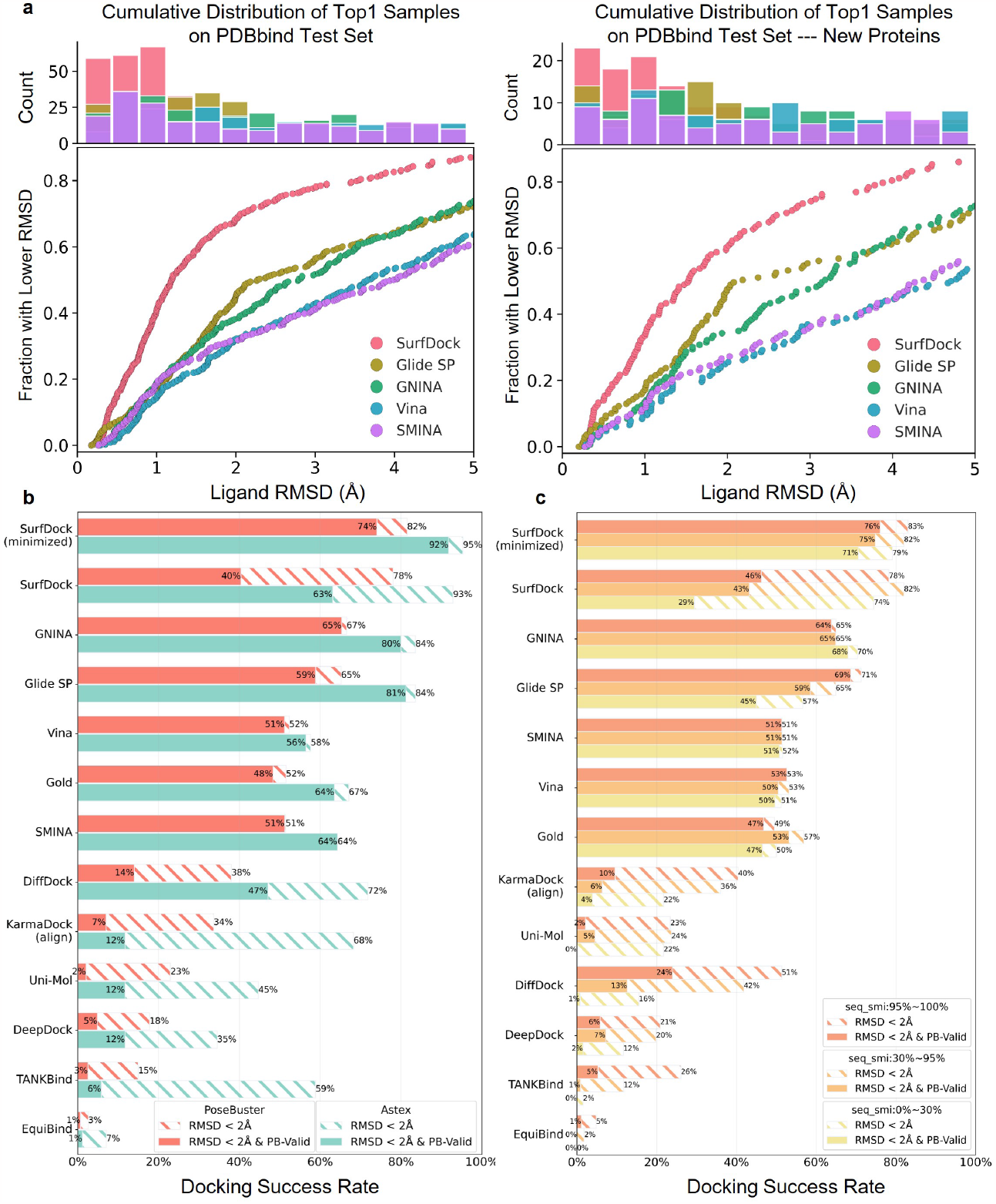
Comparative Performance of Docking Methods Across Benchmarks. The results for EquiBind, TANKBind, DiffDock, and Uni-dock are derived from existing literature, while KarmaDock is implemented from its open-sourced model weights. Glide SP, GNINA, SMINA, Vina and our SurfDock are self-implemented (details in Methods). **a**: SurfDock and traditional method performances on the PDBbind2020 time-split test set (*left*) and new proteins (*right*). Mean values are reported from three runs per method. Deep learning method comparisons are omitted due to lack of raw data. **b**: Docking method efficacy comparison using the Astex Diverse set (85 cases) as an easy test set and the PoseBusters Benchmark set (428 cases) as a challenging set. Striped bars indicate the proportion of predictions with RMSD within 2 Å; solid bars represent predictions also passing PoseBuster tests (PB-Valid), i.e., retaining biophysical restraints. **c**: Performance of docking methods on the PoseBusters Benchmark set, categorized by sequence similarity to the PDBbind2020. Striped bars show predictions with RMSD within 2 Å; solid bars denote those also PB-Valid.

**Fig. 3:**
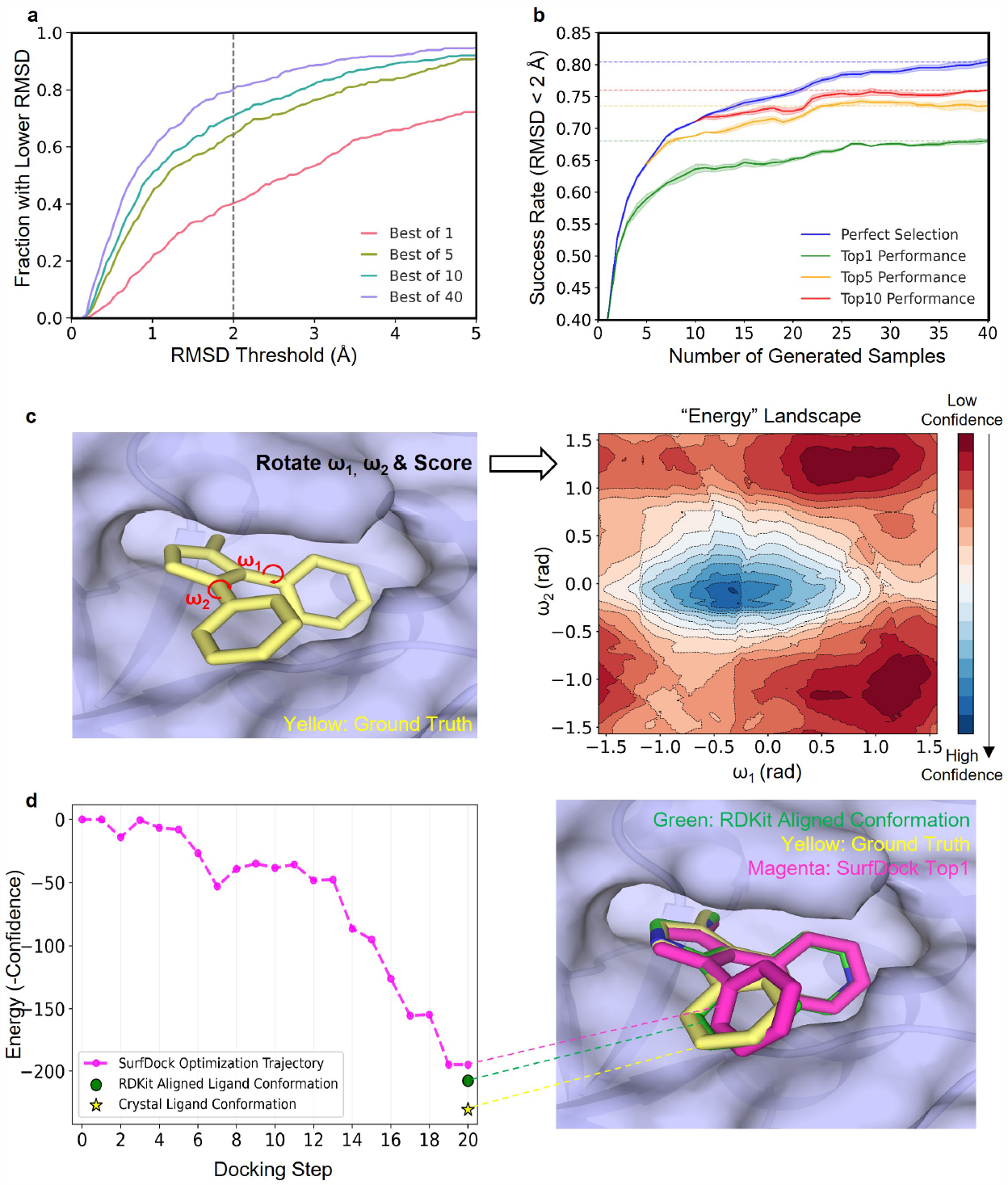
Evaluation of the Sampling Efficiency of SurfDock, the ranking ability of the scoring module SurfScore, and their consistency. **a:** Sampling Efficiency of SurfDock: This section illustrates the relationship between the number of samples and docking success rates. As the sampling number increases, there’s a corresponding increase in the likelihood of achieving success within a specified RMSD threshold. Notably, with as few as 10 samples, SurfDock demonstrates adequate efficiency. This result is averaged over three repeats. **b**: Efficacy of SurfScore. The term ‘Perfect Selection’ refers to the ideal scenario where the sample with the lowest RMSD is chosen. Remarkably, selecting the top pose from a set of 40 samples yields a 68% success rate, highlighting SurfScore’s robustness in enhancing docking precision. **c**: Torsional Profile Analysis: a specific case is presented where the scores related to the torsional profile of a ligand with two rotatable bonds are like an energy landscape. **d**: Docking as an Optimization Process: a case study where the docking procedure complemented with score estimation is analogized to a geometry optimization process. The RDKit Aligned Ligand Conformation is the RDKit generated conformation that align with the Crystal Ligand Pose, and is served as the training objective in our diffusion network.

The full end-to-end pipeline of ligand docking with SurfDock encapsulates docking, optional post-docking energy minimization, and scoring. Initially, the model identifies the protein binding pocket and initializes a user-defined number of random ligand conformations generated by RDKit^50^ from input 2D molecular graph or SMILES (Simplified molecular-input line-entry system). These random poses are then refined through a reverse diffusion process to yield final poses. If energy minimization is selected here, all poses undergo further refinement conditioned on the protein structures. Finally, SurfScore assigns a confidence score to each pose, and they are ranked accordingly. The docking-minimization-score pipeline offers a reliable system for generating ranked docking poses. This minimization stage can also be added to refine only the Top N samples selected by SurfScore for practical consideration. In our experiments, SurfDock, even without the post-docking minimization stage, achieves state-of-the-art docking success rates, underscoring its robustness and accuracy. The optional minimization stage serves to further enhance ligand validity, augmenting an already superior performance. Details of our model are provided in **Methods**.

### SurfDock Reaches State-Of-The-Art Docking Performance on Several Public Benchmark Sets

To demonstrate the effectiveness of our method, we initially selected the PDBbind2020 time-split dataset as a benchmark due to its stringent standards. In this dataset, molecules are carefully segregated to ensure no overlap between training and testing sets, thus effectively avoiding data leakage issues. This dataset features a wide spectrum of molecules, including peptides and small molecules, providing a comprehensive platform for evaluating docking capabilities. As shown in **Table 1**, SurfDock achieves a notable docking success rate (RMSD <= 2Å) of 68.41%, considerably outperforming other deep learning and traditional docking models. Additionally, when assessing docking results with RMSD under 1Å, SurfDock’s performance remains substantially superior under this rigorous metric. This advantage can be seen clearly in **Fig. 2 a**, where SurfDock clearly have more samples close to smaller RMSD when compared with the traditional docking methods we tested ourselves. To our surprise, when separating out the new proteins in PDBbind2020 test set that our model has never seen, SurfDock can still outperform all methods when comparing the metrics of Top1 samples. This separate set exhibits no ‘hard overlap’^51^ with the proteins in the training set, which means they do not possess identical structures. This indicate that the incorporation of multimodal information and diffusion generative modelling with SurfDock substantially improve the generalizability and docking success rates. We further test the rationality of generated poses using PoseBuster tool. As shown in **Table 1**, SurfDock is comparable with traditional methods in pose plausibility. If equipped with the post-docking minimization stage, the plausibility of SurfDock generated sample can gain around 19% improvements, while keeping the high docking success rate. We also compare different minimization strategies and the sequential validity check results by the PoseBusters tool in **Supplementary Table 1** and **Supplementary Fig. 1**. We show in **Supplementary Table 1** that both the docking-minimize-scoring or the docking-scoring-minimize pipeline can improve ligand validity. Here we present the docking-minimize-scoring results in **Table 1** as SurfDock(minimized).

**Table 1:** Comparative Analysis of Docking Performances on PDBbind2020 Dataset. This table presents a detailed comparison of various docking methods on the PDBbind2020 time-split test set and against novel protein targets. The results for EquiBind, TANKBind, DiffDock, E3Bind, and Uni-dock are derived from existing literature^32^, while KarmaDock’s performance is from its original publication^26^. Glide SP, GNINA, SMINA, Vina and our SurfDock are self-implemented (details in Methods). SurfDock(minimized) adopts additional post-docking minimization. Metrics include Top1/5-RMSD < 1Å/2Å and median RMSD values, with each method tested three times. We also report PB-valid (ligand poses pass all PoseBusters tests) metric for self-implemented methods. Due to the unavailability of raw data for the adopted methods, PB-valid analysis could not be conducted for them. Best results are in bold and second best are underlined in two categories.

| Performance on PDBbind2020 time-split test set (363 complexes) |  |  |  |  |  |  |  |  |  |
| --- | --- | --- | --- | --- | --- | --- | --- | --- | --- |
| Model type | Pocket | Method | Top1-RMSD |  |  |  | Top5-RMSD |  |  |
|  |  |  | %<1Å | %<2Å | Med | %<2Å & PB-valid | %<1Å | %<2Å | Med |
| DL | Blind | EquiBind | / | 5.5±1.2 | 6.2±0.3 | / | / | / | / |
| DL | Blind | TANKBind | 2.66±0.26 | 18.18±0.6 | 4.2±0.05 | / | 4.13±0.0 | 20.39±0.45 | 3.5±0.04 |
| DL | Blind | DiffDock | 15.41±0.49 | 36.62±0.35 | 3.31±0.03 | / | 21.58±0.38 | 44.19±0.49 | 2.37±0.06 |
| DL | Blind | E3Bind | / | 25.6 | 7.2 | / | / | / | / |
| DL | Given | KarmaDock | / | 56.2 | / | / | / | / | / |
| classical | Fpocket | Uni-dock | 13.33±0.4 | 18.7±0.13 | 13.2±0.26 | / | 19.16±0.39 | 27.32±0.69 | 8.3±0.25 |
| classical | P2Rank | Uni-dock | 19.31±1.07 | 28.6±1.17 | 6.40±0.22 | / | 27.76±1.03 | 39.18±1.03 | 3.76±0.06 |
| classical | PointSite | Uni-dock | 21.36±1.65 | 32.12±0.93 | 5.54±0.46 | / | 31.38±0.86 | 46.06±0.69 | 2.52±0.18 |
| classical | DiffDock | Uni-dock | 25.49±0.60 | 38.93±0.23 | 4.14±0.07 | / | 36.97±1.05 | 51.07±1.06 | 1.93±0.12 |
| classical | Given | Uni-dock | 32.77±0.38 | 51.11±0.6 | 1.89±0.04 | / | 47.5±0.23 | 67.59±0.94 | 1.11±0.02 |
| classical | Given | Glide SP | 17.36±0.00 | 44.63±0.00 | 2.27±0.00 | 38.57±0.00 | 31.13±0.00 | 60.06±0.00 | 1.54±0.00 |
| classical | Given | GNINA | 21.12±0.26 | 43.62±1.06 | 2.45±0.07 | <u>41.41±1.13</u> | 28.47±0.57 | 58.13±0.81 | 1.65±0.02 |
| classical | Given | SMINA | 18.73±0.00 | 31.68±0.00 | 3.99±0.00 | 28.37±0.00 | 28.47±0.56 | 48.48±0.00 | 2.07±0.00 |
| classical | Given | Vina | 18.32±0.02 | 36.64±0.05 | 3.42±0.01 | 32.87±0.91 | 24.79±0.00 | 50.96±0.00 | 1.87±0.01 |
| DL | Given | SurfDock | <u>40.96±0.34</u> | **68.41±0.26** | <u>1.18±0.00</u> | 36.46±0.26 | <u>54.18±0.13</u> | **75.11±0.13** | <u>0.94±0.00</u> |
| DL | Given | SurfDock (minimized) | **46.01±0.67** | <u>68.04±0.22</u> | **1.10±0.01** | **55.00±0.13** | **55.83±0.13** | <u>73.55±0.60</u> | **0.86±0.00** |
| Performance on unseen proteins in PDBbind2020 time-split test set (144 complexes) |  |  |  |  |  |  |  |  |  |
| classical | Given | Glide SP | 16.67±0.0 | 46.53±0.0 | 2.13±0.00 | 35.42±0.00 | 31.25±0.0 | 56.25±0.0 | 1.50±0.00 |
| classical | Given | GNINA | 16.67±0.0 | 38.43±1.31 | 2.75±0.18 | <u>36.57±0.87</u> | 24.31±0.57 | 53.24±1.18 | 1.82±0.06 |
| classical | Given | SMINA | 11.81±0.0 | 27.08±0.0 | 4.32±0.00 | 24.31±0.00 | 19.44±0.00 | 45.14±0.0 | 2.32±0.00 |
| classical | Given | Vina | 10.41±0.00 | 25.69±0.08 | 4.25±0.02 | 23.61±0.98 | 18.06±0.00 | 42.36±0.00 | 2.41±0.03 |
| DL | Given | SurfDock | <u>32.87±0.65</u> | <u>60.88±0.33</u> | <u>1.51±0.01</u> | 30.79±0.33 | <u>46.53±0.00</u> | <u>70.60±0.33</u> | **1.10±0.00** |
| DL | Given | SurfDock (minimized) | **37.73±0.87** | **62.50±0.57** | **1.47±0.02** | **43.75±0.56** | **47.22±0.00** | **71.06±1.31** | <u>1.12±0.00</u> |

To assess SurfDock’s efficacy more comprehensively with drug-like small molecules, we conducted evaluations using both the PoseBusters benchmark set and the Astex Diverse set, as shown in **Fig. 2 b**. These tests compared the plausibility and generalizability of generated poses across various methods. Notably, the PoseBusters benchmark set includes 428 drug-like molecule complexes released post-2021. Given that common DL docking models trained on the PDBbind2020 dataset have not been exposed to these samples, this set provides a fair basis for method comparison. The Astex Diverse set, however, is a relatively easy set, published in 2007, where most samples have been seen in the PDBbind2020 training set. In both datasets, SurfDock significantly leads in docking performance, achieving a success rate (hatched bars in **Fig. 2**) of 78% on PoseBusters set and 93% on Astex Diverse set. Compared with the other DL methods, SurfDock excels in both docking success rate and ligand validity. After the addition of post-docking minimization, the performance is further enhanced in both success rate and validity (solid bars in **Fig. 2 b**), outperforming all other DL and traditional docking methods. We also provide the cumulative distribution of top1 samples produced by different methods in **Supplementary Fig. 2 a**. We can see that SurfDock consistently outperform other methods either under RMSD<1Å or RMSD<2Å, with Glide SP and GNINA following the lead. **Supplementary Fig. 2 b and c** presents additional results including different versions of KarmaDock for a clear comparison between all competing DL methods.

Further, we evaluated SurfDock on the PoseBuster set categorized by protein sequence similarity to the PDBbind2020, as illustrated in **Fig. 2 c**. The group with low similarity can be seen as having no ‘soft overlap’^51^ with the proteins in the training set. Here, we observed that, except for SurfDock, all other DL methods exhibited significantly reduced effectiveness on proteins with less than 30% sequence similarity, regardless of pose validity. Conversely, SurfDock’s performance exhibited only a marginal decrease from familiar proteins to unfamiliar proteins in terms of docking success rate. With the enhancement of post-docking minimization, the performance of SurfDock has surpassed both DL and traditional methods on these benchmarks. SurfDock’s consistent performance across proteins with low sequence similarity highlights its exceptional ability to generalize to novel proteins. This is a critical advantage, especially considering the frequent encounter of unfamiliar protein targets in practical virtual screening tasks. The robustness and adaptability demonstrated by SurfDock not only emphasize its reliability but also its potential as a valuable tool in practical virtual screening tasks, where accurately identifying suitable ligands to novel protein targets is crucial. Considering the exceptional performance of SurfDock with the addition of minimization stage for generating accurate and reliable ligand poses, we conducted the following experiments with the minimization stage. When mentioning “SurfDock” in the following experiments, we are referring to the SurfDock with “docking-minimize-scoring” strategy unless otherwise noted.

### Evaluation of the Sampling Efficiency, Pose Selection Ability of SurfDock, and the Synergy between the Pose Generation and Scoring Module

As we have emphasized before, the effectiveness of a docking program is relied on two stages: the conformational sampling stage and the scoring stage. Accordingly, we conducted an evaluation of SurfDock’s sampling efficiency and SurfScore’s scoring accuracy independently, utilizing the PDBbind2020 time-split test set.

To discern the impact of sampling quantity on overall performance, we analyzed outcomes across varying sampling counts. Specifically, we considered a sampling effort successful if at least one instance fell within a predetermined RMSD threshold. As delineated in **Fig. 3 a**, when the sampling count reaches 10, we observe a slower rate of performance improvement with additional sampling. This indicates that SurfDock can identify a near-native ligand conformation with as few as ten samplings.

Further, we assessed the efficacy of our scoring module, SurfScore. **Fig. 3 b** illustrates that SurfScore significantly bolsters SurfDock’s performance. For instance, a single sample per ligand yields a docking success rate of around 40%. However, generating 40 samples and applying SurfScore to select the top pose elevates the success rate to over 65%. While there remains a disparity between this outcome and ‘perfect selection’ – the ideal scenario of ranking the most accurate ligand pose at the top from all samples – SurfScore’s current capability suffices for practical applications.

To better illustrate that SurfScore captures key interactions between proteins and ligands, we present a specific case in **Fig. 3 c**. Here, a ligand with two rotatable bonds is analyzed. By treating the crystal ligand pose as a reference point and varying the torsional angles ω_1_ and ω_2_, we observe the scoring trends from SurfScore. Interpreting these scores as energy values reveals a landscape centered around the reference pose, with a plausible distribution of local minima as torsional angles shift. Building upon this, we explored the consistency between our docking and scoring modules, as they share the same representational framework and are separately trained on the same crystal protein-ligand complex data. **Fig. 3 d** showcases a sequential record of docking outputs and their corresponding SurfScore evaluations. In the dynamic progression of the docking process facilitated by SurfDock, there is a notable trend where the generated ligand poses incrementally gravitate towards lower energy states (or higher confidence). This evolution often involves navigating through and overcoming local energy minima, ultimately resulting in an alignment that is increasingly proximate to both the RDKit aligned pose and the crystal ligand pose. It is important to clarify that the RDKit aligned pose refers to a conformation generated by RDKit aligned to the crystal ligand pose, and is utilized as training objective for our diffusion network, as explained in **Methods**. This aligned pose can be regarded as a ‘theoretical limit’ for the generation module of SurfDock in the absence of additional refinements. However, with the integration of our post-docking minimization strategy, SurfDock demonstrates the potential to identify ligand poses that surpass the RDKit aligned pose in terms of energies estimated by our scoring module. We have included several such examples in **Supplementary Fig. 4**.

These findings highlight the effective synergy between the docking and scoring processes, demonstrating their combined strength in capturing crucial protein-ligand interactions during generative modeling. The high degree of consistency between the two modules, despite their separate training phases, can be attributed to their aligned objective of learning the distribution of crystal structures, which likely plays a key role in their harmonized performance.

### Influence of ligand flexibility on docking performance

In molecular docking, ligand flexibility critically influences conformation sampling efficiency^52^. This relationship becomes increasingly complex as the number of rotatable bonds and heavy atoms in the ligand rises, expanding the search space for potential conformations. We first count the distribution of the number of rotatable bonds and heavy atoms on PDBbind2020 time-split test set in **Supplementary Fig. 3**. We find that the distribution is quite large, ranging from 0 to 75 for the number of rotatable bonds, or 6 to 150 for the number of heavy atoms. Thus, the ligand flexibility in this dataset is challenging for both DL and traditional docking methods. Our experimental results, depicted in **Fig. 4 a** and **b**, corroborate this trend, aligning with findings^52^ by Hou et al. We observed a significant decline in the performance of traditional docking methods when ligands possess near or more than 15 rotatable bonds, or approximately 35 heavy atoms, on the PDBbind2020 time-split test set. SurfDock, however, demonstrates notable proficiency in handling ligands within these ranges, often matching or surpassing traditional methods. On the other hand, it is widely acknowledged that the majority of drugs and drug-like compounds typically contain fewer than 10 rotatable bonds^52^. Within this subset, SurfDock’s performance is particularly striking, achieving an efficacy rate close to 80%, which represents a substantial improvement of approximately 20% over conventional methods.

**Fig. 4:**
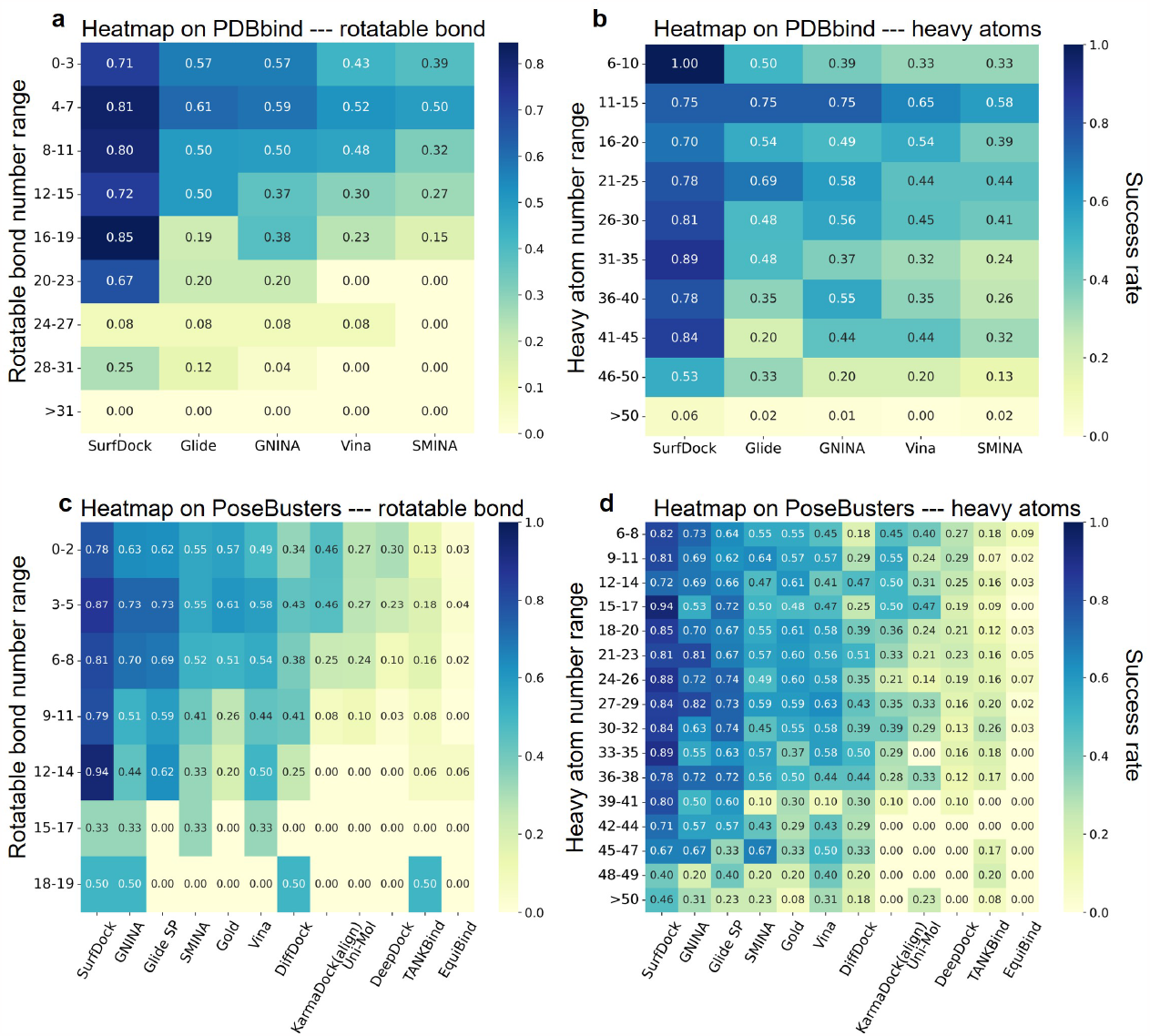
The Performance across Different Docking Methods on PDBbind2020 time-split test set and PoseBusters Benchmark set with the number of rotatable bonds and heavy atoms. **a, c:** Impact of the number of rotatable bonds on docking accuracy. **b, d:** Impact of the number of heavy atoms on docking accuracy.

We extended our investigation to the PoseBusters Benchmark Set, which primarily comprises drug-like molecules. This set presents a distribution of rotatable bonds and heavy atoms smaller to those in the previous dataset, also depicted in **Supplementary Fig. 3**. Consistent with expectations based on the molecular characteristics typical of drug-like compounds, SurfDock exhibits a remarkable performance across varying counts of rotatable bonds and heavy atoms, as shown in **Fig. 4 c** and **d**. This performance not only aligns with our observations from the PDBbind2020 set but also distinctly demonstrates SurfDock’s superiority or at least equivalence to traditional and other deep learning-based docking methods, especially in handling drug-like molecules.

These findings underscore SurfDock’s potential in facilitating drug discovery processes. Despite these promising results, we acknowledge the limitations of SurfDock in handling larger molecules like peptides. This constraint could stem from the scarcity of large ligand training data in PDBbind, as indicated in **Supplementary Fig. 3**. Addressing this challenge will be a focus of our future research, aiming to extend SurfDock’s applicability and efficacy in molecular docking.

### SurfDock Can Serve as A Tool for Virtual Screening with Excellent Performance

To further investigate the virtual screening capabilities of SurfDock, we conducted a preliminary evaluation of SurfDock’s virtual screening capabilities using the DEKOIS2.0 dataset^47^. This dataset, comprising both active ligands and inactive decoys, includes 81 varied targets. Each target is associated with 40 active compounds and 1,200 inactive decoys. This diverse and challenging benchmark set serves as an ideal platform to test the efficacy of SurfDock in discerning active ligands from decoys.

Considering that efficiency is important in practical virtual screening task, here we adopt the “docking-scoring-minimize-rescoring” approach. In particular, we first generate 40 samples and select the Top 10 samples. We further minimize the 10 selected samples and re-score them using another version of SurfScore that is specifically trained for virtual screening task for fair comparison with other methods, as detailed in **Methods**. Finally, the Top 1 sample is used for evaluation. The results, illustrated in **Fig. 5**, position SurfDock at the forefront of current docking algorithms in terms of performance. A notable highlight is SurfDock’s achievement in the metric EF 0.5%, reaching 21.00. This is significant in virtual screening, especially when dealing with large libraries of compounds. The primary goal of a docking algorithm in this context is to prioritize or ‘enrich’ the subset of compounds that are most likely to be active, thus reducing the number of compounds that need to be further tested in more resource-intensive experiments. The efficacy of SurfDock in identifying active candidates at the top of the list is critical in large-scale virtual screening processes. This success is in line with prior benchmarks that attest to SurfDock’s ability to generate accurate and reliable ligand poses. In contrast, although KarmaDock may generate less plausible poses, it surprisingly outperforms established methods like Glide SP in virtual screening tasks. As reported in the original KarmaDock publication, the algorithm’s other two versions, despite having lower plausibility, also demonstrate effective performance on the DEKOIS2.0 dataset. These results highlight the necessity for more in-depth research and stringent benchmarking to understand the factors influencing the efficacy of docking algorithms in virtual screening.

**Fig. 5:**
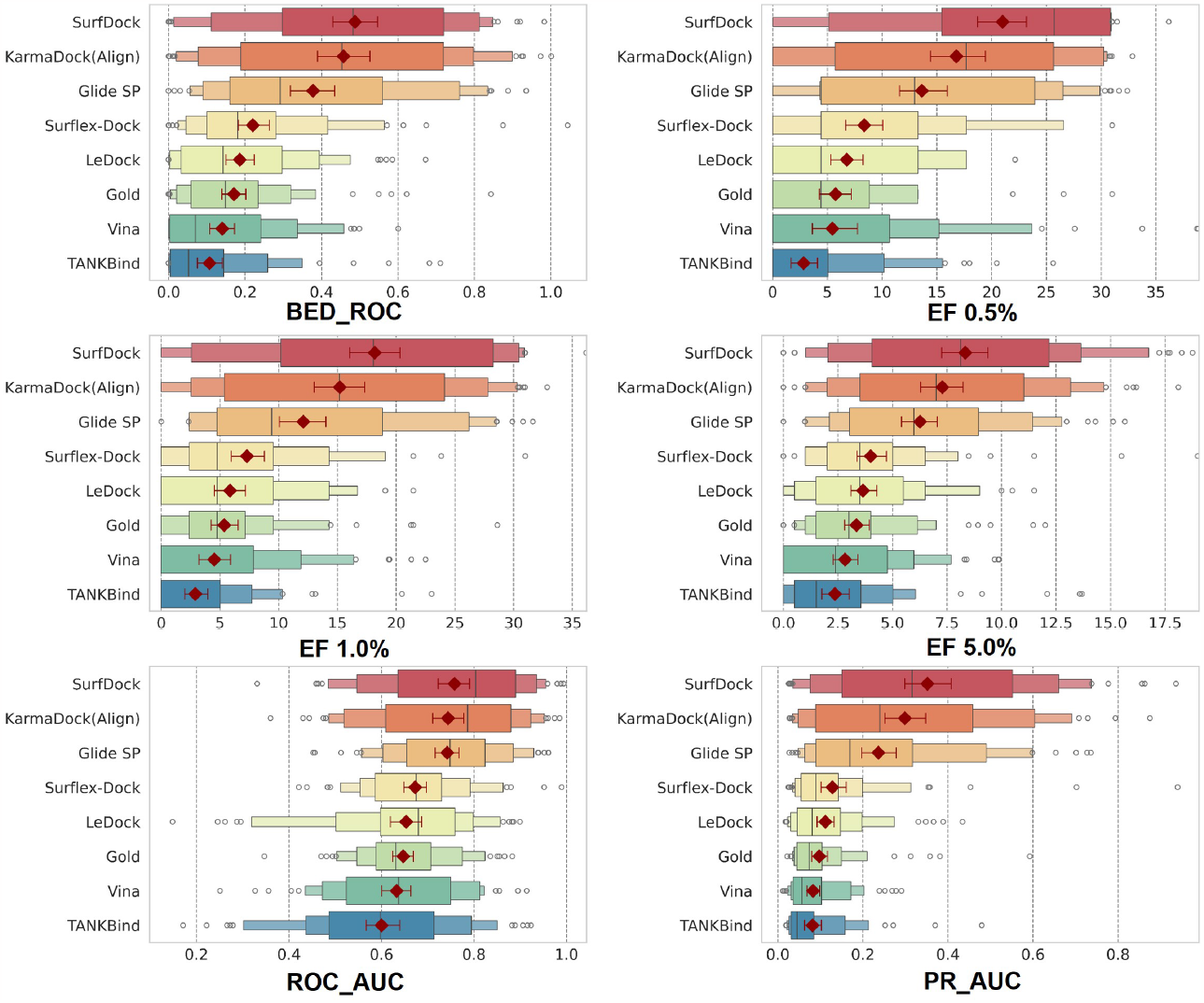
The Performance of Different Docking Methods on DEKOIS2.0 Dataset. The results except SurfDock are adopted from the publication of KarmaDock^26^. The boxenplot illustrates the distribution of key metrics for each model, highlighting data spread and variability. Superimposed red diamonds represent the mean values. This figure displays the performance comparison of various methods on the DEKOIS2.0 dataset, featuring key metrics: Boltzmann-Enhanced Discrimination of Receiver Operating Characteristic (BED_ROC), which focuses on the early identification of active compounds; Enrichment Factors (EF) is defined as the percentage of active ligands observed among all of the active ligands for a given percentile of the top-ranked candidates (0.5%, 1.0%, or 5.0%) of a chemical library; Receiver Operating Characteristic Area Under the Curve (ROC_AUC), assessing overall classification accuracy; and Precision-Recall Area Under the Curve (PR_AUC), evaluating the trade-off between precision and recall, particularly in datasets with class imbalances.

Next, to evaluate the scoring efficacy of our module, SurfScore, we utilized poses generated by traditional methods, and then reassessed their binding affinities using SurfScore. The outcomes of this assessment are presented in **Supplementary Fig. 5**. The combination of Glide SP/Surflex-Dock with SurfScore shows comparable results to SurfDock across all evaluation metrics, although SurfDock maintains a lead in the EF 0.5% metric. This observation indicates that Glide SP and Surflex-Dock exhibit robust sampling capabilities on the DEKOIS2.0 benchmark set. This is consistent with earlier research highlighting the effectiveness of both Glide SP and Surflex-Dock in accurately sampling conformations^52^. Additionally, our previous experiments, as illustrated in the PoseBusters Benchmark Set (**Fig. 2 b**) and **Supplementary Fig. 2 a**, affirm Glide SP’s strength as a docking algorithm, especially in benchmarks with simpler ligand compositions. Support for this comes from **Supplementary Fig. 3**, which reveals that most ligands and decoys in DEKOIS2.0 have fewer than 20 rotatable bonds. Thus, the observation that SurfDock and Glide SP performs similar in sampling power is plausible. It’s important to note that in this virtual screening experiment, we chose the less accurate “docking-scoring-minimize-rescoring” for a more efficient testing setup. We anticipate that by optimizing our workflow, SurfDock’s performance can be further enhanced in practical virtual screening tasks.

## CONCLUSION

In this Article, we have introduced SurfDock, an advanced geometric diffusion network tailored for generating reliable binding ligand poses conditioned on protein pockets and ligand 2D graph or SMILES. SurfDock also integrates a comprehensive internal scoring module, SurfScore, for confidence estimation, suitable for virtual screening tasks.

Throughout our research, SurfDock has demonstrated exceptional performance across various benchmarks, including PDBbind2020 time-split set, the Astex Diverse Set, and the PoseBusters Benchmark set. Its ability to integrate multimodal protein information—encompassing surface features, residue structure, and pre-trained sequence-level features—into a cohesive surface node level representation has been instrumental in achieving high docking success rates and improved plausibility. Another aspect of SurfDock’s functionality is its optional force field relaxation step, designed for protein-fixed ligand optimization, which significantly enhances its accuracy. This feature, along with pose generation and scoring, allows SurfDock to outperform existing DL and traditional methods in both docking success rates and pose rationality. More importantly, SurfDock demonstrates remarkable adaptability to new proteins and is highly effective for practical virtual screening. Its combination of strong performance and practical utility highlights its considerable promise in SBDD.

In summary, we have shown that diffusion generative modeling, enhanced with multi-modal information, excels in pocket-aware ligand docking, surpassing traditional docking and DL methods. This makes SurfDock a valuable asset to the SBDD community, offering new avenues for drug discovery and protein-ligand interaction studies.

## METHODS

### Overview

Our model, SurfDock, comprises two main components: a docking module and a scoring module. Both modules receive input from a multimodal feature fusion layer that integrates sequence, structure, and surface features. SurfDock is built upon an *E*(3)-equivariant, diffusion-based graph neural network, while the scoring module is constructed from an equivariant graph neural network paired with an invariant mixture density network.

The challenge in developing deep learning models for molecular docking arises from the inherent aleatoric uncertainty related to pose prediction, where multiple poses could be correct, and the epistemic uncertainty stemming from the complex nature of the task relative to the limited model capacity and available data^27^. Despite advances in cryo-electron microscopy and crystallography, high-quality protein-ligand complex data remains scarce, necessitating an architecture that can generalize well with limited high-quality structural information. Research indicates that equivariant networks are highly data-efficient, achieving superior performance with less data^53^, which makes it a great choice for our situation.

Besides equivariant neural networks, we introduce surface information upon to a diffusion model. The molecular surface is a higher-level representation of protein structure, modeling a protein as a continuous shape with geometric and chemical features. This information allows the diffusion model to better perceive the protein’s surface geometry, potentially avoiding physically improbable pose predictions too close to protein atoms.

Our model, an *E*(3)-equivariant, diffusion-based graph neural network, follows the generative model training paradigm and is well-suited for molecular docking, a task characterized by limited data but high complexity. Unlike methods that represent proteins and ligands at the atomic level and predict coordinates for each atom, SurfDock is trained through a process that incrementally distorts the native conformation at various degrees, enabling the model to learn how to restore the correct conformation. In docking, bond lengths and angles can be swiftly and accurately determined by standard cheminformatics methods^54^. We consider only the torsion degrees of freedom m, where m is the number of torsion angles, and six degrees of freedom for translation and rotation, significantly narrowing the problem scope. SurfDock takes a seed conformation *c* ∈ ℝ^3×*n*^ of the ligand as input and alters only the relative position and torsion degrees of freedom in the final bound conformation. Thus, our problem is defined on an (*m* + 6) -dimensional submanifold *M*_*c*_ ⊂ ℝ^3×*n*^, formulating molecular docking as learning a probability distribution *p*_*c*_ (x|y) over the manifold, over the manifold, conditioned on a protein pocket structure y.

Finally, we follow a similar approach to DiffDock in training the diffusion model on the product space of three subspaces: ligand rotation, translation, and torsion. The input to our model is the crystal conformation of the protein pocket and a seed conformation of the ligand. The output comprises *m* scalar torsion angles and two translation-rotation vectors for each ligand. Following docking, the SurfScore module receives the docked complex and outputs a scalar score for the complex

### Details of feature processing

In the feature processing methodology for SurfDock, a geometric heterogeneous graph is constructed, incorporating ligand, receptor residues, and surface nodes. The interactions among these components are defined with specific cutoffs and interaction rules:

#### Ligand Atoms-Ligand Atoms Interactions

These interactions are defined using a 5Å cutoff, aligning with standard practices for atomic interactions. Covalent bonds are additionally preserved as separate edges, providing detailed chemical structure information.

#### Receptor Residues-Receptor Residues Interactions

For interactions between receptor residues, a cutoff of 15Å is used, with a maximum of 30 neighbors allowed for each residue. This approach helps to capture significant inter-residue interactions while maintaining computational efficiency.

#### Receptor Residues-Surface Nodes Interactions

For interactions between receptor residues and surface nodes, a cutoff of 15Å is used, with a maximum of 30 neighbors allowed for each surface node to maintain computational efficiency.

#### Surface Nodes-Surface Nodes Interactions

Similar to DeepDock, each edge 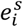 is represented by a vector indicating the relative Cartesian coordinates of the connected nodes, providing spatial context for these interactions.

#### Surface Nodes-Ligand Atoms Interactions

These interactions use a cutoff of 10 + 3σ_*tr*_ Å, where σ_*tr*_ represents the current standard deviation of the translational diffusion noise. This dynamically adjusts the interaction range based on the uncertainty in the diffusion process, ensuring high-probability interactions in the final pose are included in the message passing at every step.

The ligand in SurfDock is represented as an attributed graph *G*^*l*^ = (*V*^*l*^, *E*^*l*^), with *V*^*l*^ representing atoms and *E*^*l*^ representing edges. Ligand atom features include atomic number, chirality, degree, formal charge, implicit valence, number of connected hydrogens, number of radical electrons, hybridization type, aromaticity, ring membership, and ring size (3 to 8). These features are enriched with sinusoidal embeddings of diffusion time. Edge features include bond type, ring status, conjugation, stereochemistry, and radial basis embeddings of edge length.

The protein residue graph is denoted *G*^*α*^ = (*V*^*α*^, *E*^*α*^), with each node representing a residue at the C_*α*_ position. Node features include amino acid type, language model embeddings from ESM-2, and features used in the RTMScore model. Edge features are informed by RTMScore and include radial basis embeddings of edge length.

Surface generation follows the DeepDock and MaSIF process. Surfaces are triangulated using MSMS, with specifications of density and probe radius as per MaSIF guidelines, and processed using PyMesh^55^. The resulting mesh *G*^*s*^ = (*V*^*s*^, *E*^*s*^) comprises nodes 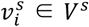 and edges 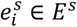 . Node features include Poisson– Boltzmann electrostatics, free electrons and proton donors, hydropathy, shape index, and sinusoidal embeddings of diffusion time. Edge features are defined by relative Cartesian coordinates (vector) and radial basis embeddings of edge length (scalar).

Scalar features of each node and edge are transformed using learnable two-layer MLPs into a set of scalar features for initial representations in the interaction layers. Only nodes defining the binding site (within 8Å of any ligand atom) are used to train the model, focusing on the most relevant interaction sites.

### Model architecture

The docking module in SurfDock is an advanced *E*(3)-equivariant, diffusion-based graph neural network that utilizes tensor products of irreducible representations (irreps), following the conventions defined in the e3nn library^56^. This framework effectively incorporates both equivariant and invariant features for robust representation learning.

### Residue-residue intra-interaction

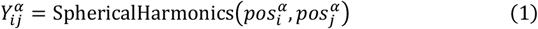

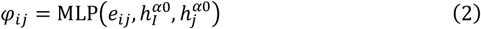

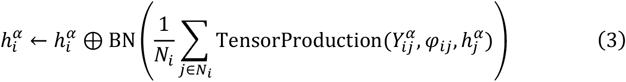

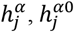 represent the residue’s features and initial scalar features, respectively. 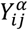 are the spherical harmonics computed up to *l* = 2, and BN denotes batch normalization. The output orders in this process are restricted to a maximum of *l* = 1.

### Residue-surface inter-interaction

In the residue-surface inter-interaction layer of SurfDock, the updated residue node representations are further integrated with surface node information. Once the connected graph structure is established, node messages are updated via the Tensor Product Layer, following a sequence of operations:

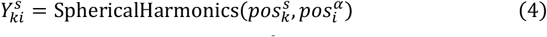

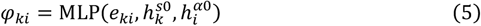

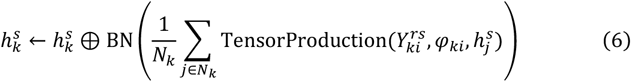

This module mirrors the earlier one in function but differs in the types of nodes and edges involved in the convolution process.

### Surface-ligand inter-interaction

In the surface-ligand inter-interaction stage of SurfDock, both the ligand and surface undergo internal updates similar to the residue-residue intra-interaction process. This step involves updating the ligand and surface using a consistent architecture, yielding new representations: 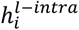 for the ligand and 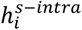 for the surface. Concurrently, akin to the residue-surface interaction layer, we construct a ligand-surface radius graph to facilitate information exchange between the ligand and surface, generating representations: 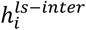for ligand-to-surface and 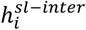 for surface-to-ligand interactions. The final representations of the ligand and surface in SurfDock are updated through an integration of inter- and intra-interaction features, as per the following equations:

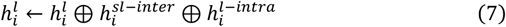

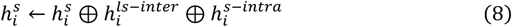

Following the final interaction layer, the updated ligand node representations are employed to generate the outputs. To predict the translation a a convolution operation is performed on each ligand atom with the unweighted center of mass *c*. This approach is in alignment with the methodology used in DiffDock, allowing for accurate determination of ligand pose in relation to the target surface:

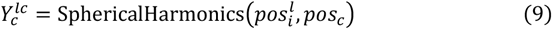

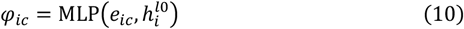

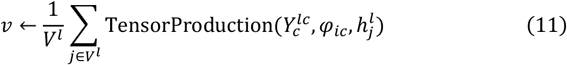

Following the strategy in DiffDock, the output *v* for ligand translation and rotation scores is constrained to include two odd parity vectors and two even parity vectors. This composition is essential in the context of the coarse-grained model used for protein representation, where the parity of the scoring output is not distinctly even or odd. Then, we integrate the even and odd components of *v*, adjusting their magnitude while preserving their original directional characteristics with an MLP. This MLP incorporates the current magnitude and the sinusoidal embeddings of the diffusion time *s*_*t*_. The following equations detail this process:

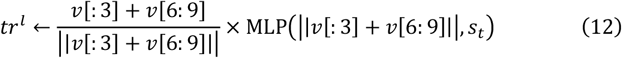

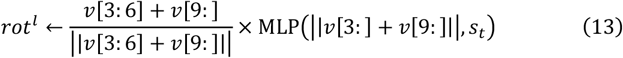

For the torsional score in SurfDock, we adopt a methodology similar to Torsional Diffusion for predicting a scalar score *δ*_*tor*_ for each rotatable bond *g* = (*g*_0_, *g*_1_). This prediction involves convolving the neighbor atoms in a radius graph with the center *p* of the bond. The convolutional filter *T*_*g*_ for each bond *g* is constructed from the tensor product of the spherical harmonics representation (with *l* = 2) of the bond axis 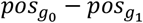, as detailed in the following steps:

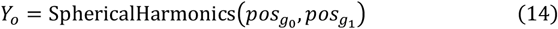

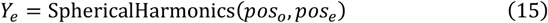

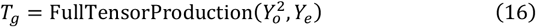

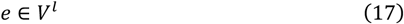

The convolutional filter *T*_*g*_ is then utilized to convolve with the representations of every neighboring atom within the radius graph, as per the following procedure:

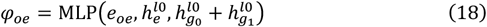

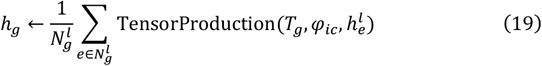

Finally, the torsional score is refined using a two-layer LP featuring a tanh nonlinearity and no biases. This MLP output is then “denormalized” by multiplying with the expected magnitude of a score in *SO*(2), adjusted by the diffusion parameter *δ*_*tor*_.

### Transformation of the ligand conformation

During each inference step in SurfDock, the ligand conformation is updated using translation, rotation, and torsion scores. The update process involves a unified global translation, where all ligand atoms are simultaneously translated and rotated around the ligand’s geometric center. However, updating the ligand torsion angles is particularly critical in the docking process. To address the potential perturbation of the ligand’s center of mass position following torsion angle updates, we employ RMSD alignment, as suggested in DiffDock. This alignment method operates by realigning the ligand, post-torsion angle adjustment, to its original pose prior to the torsion changes.

### Ranking and Screening module

In SurfDock, we introduce SurfScore, a scoring module designed to enhance pose ranking and screening capabilities. SurfScore’s input architecture mirrors that of the docking module, retaining interaction layers for residue-residue, residue-surface, surface-surface, and ligand-ligand interactions, while excluding the surface-ligand interaction layer. It employs a mixed density network (MDN) for learning the distance statistical potential between protein surface and ligands.

The process begins with extracting ligand and surface node representations 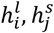, which are concatenated and fed into an MDN^49^. The MDN uses an MLP to generate a hidden representation 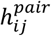 integrating both target and ligand node data. This is mathematically represented as follows:

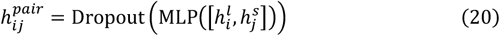

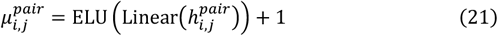

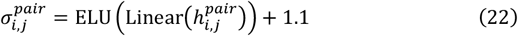

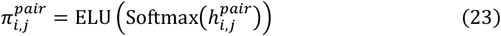

The hidden representation is used to compute the outputs of the MDN, encompassing means 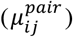, standard deviations 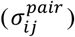 and mixing coefficients 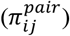. These parameters are pivotal in formulating a mixture of Gaussians. In this context, a complex mixture of 20 Gaussians models the probability density distribution pertaining to the distance between ligand and target nodes.

Further, the extracted ligand node features 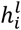 were used for predicting auxiliary tasks, specifically atom and bond types in relation to neighboring nodes. This approach is inspired by findings from DeepDock, which highlighted the benefits of auxiliary tasks in learning molecular structures, thereby expediting the training process. All MLPs used are composed of a linear layer followed by batch normalization and an Exponential Linear Unit (ELU) as activation function. A consistent dropout rate of 0.1 was maintained across experimental setups.

### Training details

#### For docking

In the docking experiments, we aligned our data and partitioning strategy with EquiBind and DiffDock, ensuring that test data comprised entirely unseen ligands. To address the distribution shift encountered during inference due to the use of RDKit-generated conformations, our training objective was reformulated to align with the conformation closest to the ground-truth pose. At each time step *t*, the input ligand pose is subject to random perturbations, which include:

#### Translational perturbation

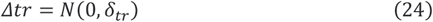

#### Rotational perturbation

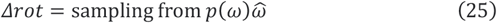

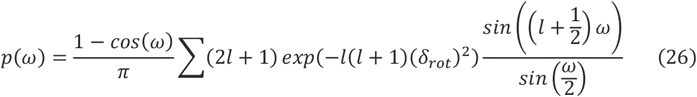

#### Torsional perturbation

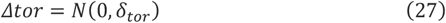

Here, *p*(*ω*) represents the isotropic Gaussian distribution on *SO*(3), and the 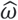 is a unit vector from random sampling. The training utilizes a score-based diffusion generative model on a Riemannian manifold, sampling and regressing against the diffusion kernel’s score. Our methodology ensures orthogonality between torsional and rot-translational updates. The training employs separate loss functions for translation (*L*_*tr*_), rotation (*L*_*rot*_) and torsion (*L*_*tor*_), with the final loss function being:

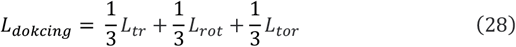

The diffusion model is trained until no further improvement is observed on the validation set within 50 epochs.

### For the Scoring Module

Different from the training of diffusion model where RMSD-aligned conformations to mitigate training-inference data drift, the scoring module directly use crystal protein-ligand complex conformation for training to learn the distance statistical distribution. The training is governed by the following equations:

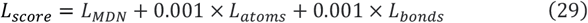

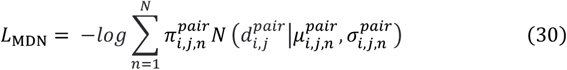

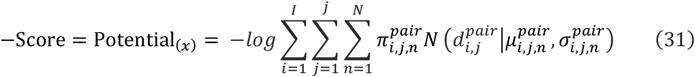

The L_*MDN*_ focuses on minimizing the negative log-likelihood of 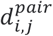, measuring the distance between surface node *v*^*s*^ and ligand node *v*^*l*^. This is computed using a mixture model composed of 20 Gaussians, parameterized by predicted 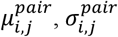 and 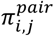. Additionally, *L*_*atoms*_ and *L*_*bonds*_, the cross-entropy cost functions for predicting atom and bond types, serve as auxiliary tasks. The L_*MDN*_ in equation (30) can be adapted to define a potential function, *Potential*_(*x*)_, specifically tailored for evaluating a given target-ligand complex. In practice, this potential function is instrumental in scoring protein-ligand complexes, enabling the assessment of the conformational rationality of compounds. It is a pivotal tool for compound screening, where the lower value of Potential_(*x*)_ (or a higher score) correlates with a higher likelihood of the target-ligand complex being in a particular conformation.

Training was conducted for 60 epochs with a batch size of 16. During training, contributions from ligand-target node pairs with 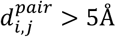 were masked. In inference, this masking threshold was adjusted to 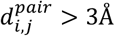.

For the virtual screening task, SurfScore was retrained using a random data split from PDBBind2020 to be comparable with other baseline models. During training, contributions from ligand-target node pairs with 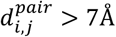 were masked. In inference, this masking threshold was adjusted to 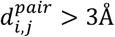..

### Post-docking energy minimization protocol

Following Deane et al.^34^, we performed post-docking energy minimization using OpenMM^57^ with AMBER ff14sb^58^ for proteins and Sage^59^ (or GAFF^60^ for incompatible ligands) for small molecules. Protein structures were prepared with PDBfixer^57^ as in Alphafold2^4^. During minimization, we fixed protein atoms, allowing only ligand atoms to move, ensuring focused energy optimization of ligands in the binding pocket.

## Supporting information

SurfDock-Supplementary-Information

## Data availability

The protein-ligand complexes of PDBBind v2020 preprocessed as described in the paper “EquiBind: Geometric Deep Learning for Drug Binding Structure Prediction” https://zenodo.org/records/6408497

The protein-ligand complexes of the Astex Diverse set and the PoseBusters Benchmark set as described in the paper “PoseBusters: AI-based docking methods fail to generate physically valid poses or generalise to novel sequences” https://zenodo.org/records/8278563

## Code availability

The code used to generate the results shown in this study is available under an MIT Licence in the repository.

Code will be available after our paper has been published at: https://github.com/CAODH/SurfDock

