## Supplementary material for "SurfDock is a Surface-Informed Diffusion Generative Model for Reliable and Accurate Protein-ligand Complex Prediction": SurfDock-Supplementary-Information

Duanhua Cao,[▽,1,2] Mingan Chen,[▽,2,3,4] Runze Zhang [▽,2,5], Jie Yu[2,3,6], Xinyu Jiang[2,5], Zhehuan Fan[2,5], Wei Zhang[2,5], Mingyue Zheng*[,1,2,5]

[1]Innovation Institute for Artificial Intelligence in Medicine of Zhejiang University, College of Pharmaceutical Sciences, Zhejiang University, Hangzhou, Zhejiang 310058, China

[2]Drug Discovery and Design Center, State Key Laboratory of Drug Research, Shanghai Institute of Materia Medica, Chinese Academy of Sciences, 555 Zuchongzhi Road, Shanghai 201203, China

[3]School of Physical Science and Technology, Shanghai Tech University, Shanghai, 201210, China

[4]Lingang Laboratory, Shanghai, 200031, China

[5]University of Chinese Academy of Sciences, No. 19A Yuquan Road, Beijing 100049, China

[6]School of Information Science and Technology, Shanghai Tech University, Shanghai, 201210, China

**Table of Contents**

S1 Docking Protocols

S2 Waterfall Plot of PoseBusters Sequential Checks

S3 Additional Results on PDBbind Test Set

S4 Additional Results on Astex and PoseBusters Benchmark Sets

S5 Distribution of Rotatable Bonds and Heavy Atoms

S6 SurfDock Optimization Trajectories

S7 Re-scoring Performance

### S1 Docking Protocols

Here we list the docking methods we implemented ourselves. The details of other methods on PDBbind test set and PoseBusters Benchmark set can be found in work[1] by Ke et.al., and work[2] by Deane et.al.

#### SurfDock

*Parameters*

Besides the parameters we mentioned in **Method**, we set diffusion steps to 20 throughout the experiments. For docking tasks, we set sampling number to 40; for the virtual screening tasks, we set sampling number to 10 for efficiency.

#### Vina

*Software version*

Vina 1.2.3 , Reduce 3.23.130521, ADFRsuite 1.0， RDKit 2022.09.1

*Ligand preparation*

The ligands are processed into PDBQT files using the ADFR *prepare_ligand* scripts and then their centroid coordinates and coordinate ranges in different directions are calculated using RDKit.

*Protein preparation*

Hydrogen atoms were added with reduce and then the PDBQT files were generated with the ADFR *prepare_receptor* script

*Parameters*

Use the center of the co-crystallized ligand along with the maximum diameters in three directions of the ligand + 4 Å to set the size of the docking box. Vina was used to create 40 poses with an exhaustiveness setting of 32 and the top-ranked pose was selected

#### SMINA

*Software version*

SMINA python API 2021.12.09

*Ligand preparation*

The generated starting ligand conformations were used without further processing

*Protein preparation*

The generated starting protein conformations were used without further processing

*Parameters*

Use the center of the co-crystallized ligand and *autobox_add* is set to 4 Å. SMINA was used to create 40 poses with an exhaustiveness setting of 32 and the top-ranked pose was selected

#### GNINA

*Software version*

Gnina 1.0.3 from authors' public code repository https://github.com/gnina/gnina.

*Ligand preparation*

The generated starting ligand conformations were used without further processing.

*Protein preparation*

The generated starting protein conformations were used without further processing.

*Parameters*

The default parameters as illustrated in https://github.com/gnina/gnina were used, with maximum number of generated poses 40.

**Glide SP**

*Software version*

Schrödinger (version 12.6; Schrödinger, LLC: New York, NY, 542 2020) was used throughout this work, with Glide module (version 8.9) using the standard precision (SP) mode.

*Ligand preparation*

LigPrep module was used with default settings.

*Protein preparation*

Protein Preparation Wizard module was used with default settings.

*Parameters*

The Receptor Grid Generation module was used with default settings to generate receptor grid centered on the co-crystallized ligand.

**KarmaDock**

*Software version*

We use KarmaDock 1.0.1 and the provided checkpoint with all default settings from authors' public code repository: https://github.com/schrojunzhang/KarmaDock.

### S2 Waterfall Plot of PoseBusters Sequential Checks

The PoseBusters tool provides the following physical and chemical checks:

1. *If the docked ligand can be successfully loaded*
2. *If RMSD is within 2 Å*
3. *If the molecule can pass RDKit sanity check*
4. *If the molecular formula is preserved compared with the crystal ligand*
5. *If the molecular bonds are preserved compared with the crystal ligand*
6. *If the sp3 stereochemistry is preserved compared with the crystal ligand*
7. *If the double bond stereochemistry is preserved compared with the crystal ligand*
8. *If bond lengths are within bounds from RDKit's Distance Geometry module*
9. *If bond angles are within bounds from RDKit's Distance Geometry module*
10. *If there exist internal clashes in the ligand*
11. *If atoms in the aromatic ring are in a plane*
12. *If atoms in the double bond are in a plane*
13. *If the energy ratio is within threshold determined by RDKit UFF*

*14. If there are clashes between ligand and protein*

*15. If there are clashes between ligand and organic cofactors*

*16. If there are clashes between ligand and inorganic cofactors*

We present the sequential check results of SurfDock, SurfDock(minimized), Glide, GNINA, SMINA on the three benchmark sets we tested in **Supplementary Fig. 1**. In the original publication of PoseBusters, the authors show that the best DL model DiffDock generates many ligand conformations having clashes with the protein, which is the main cause that reduces the success rate of DiffDock on PoseBusters benchmark set from 38% to 14%. Here, the success rate of SurfDock after considering pose validity is about 40% (173 of 428), with significant improvement over DiffDock. After adding the minimization stage, the success rate considering validity increases to about 74% (317 of 428). The minimization stage greatly reduces the number of protein-ligand clashes, which aligns with the findings in PoseBusters paper.

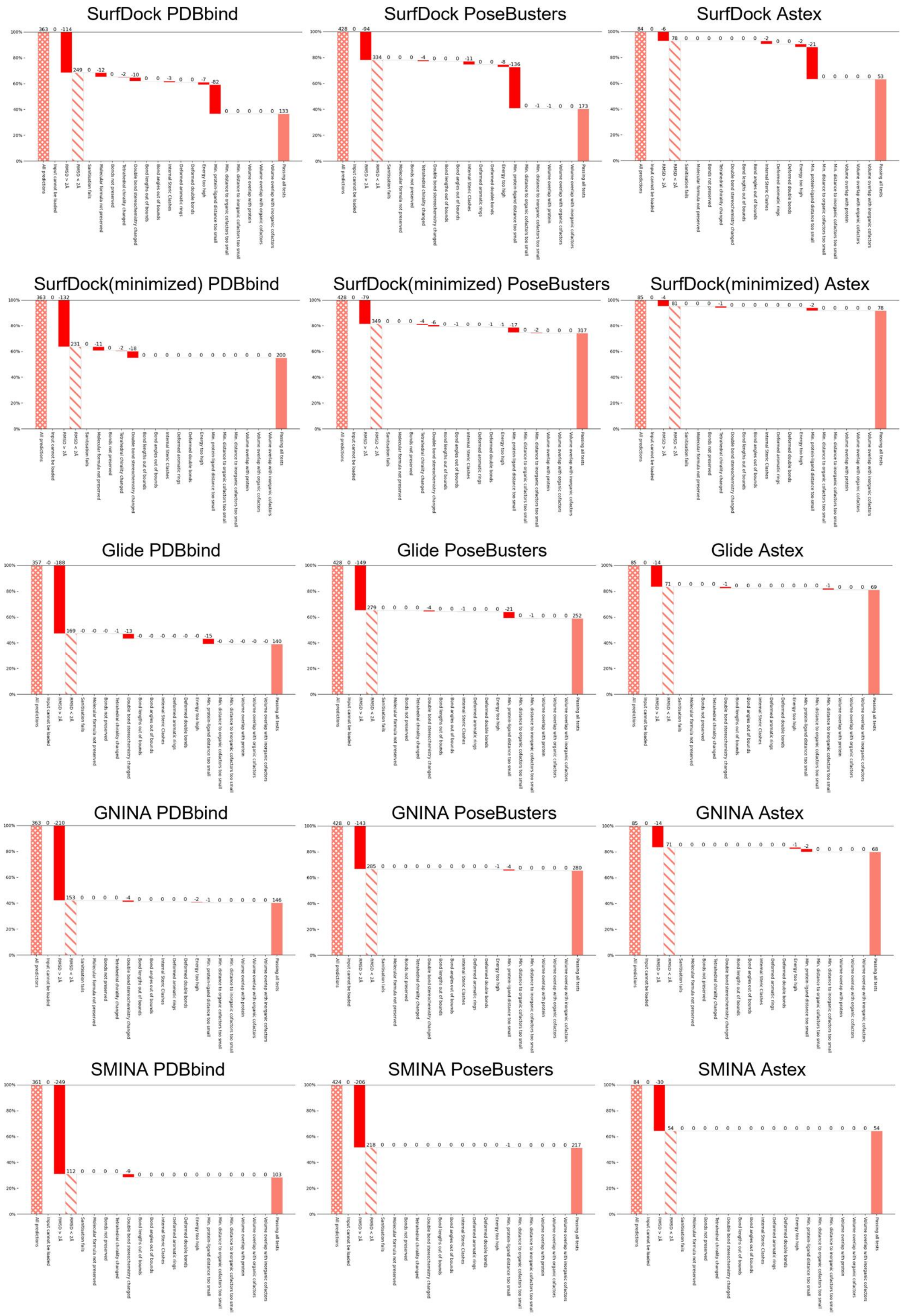


**Supplementary Fig. 1 | Sequential PoseBusters check results of different models across different datasets.** For results on PDBbind, we randomly select one repeat for each method as the plausibility checks between repeats are similar.

### S3 Additional Results on PDBbind Test Set

We compared two computational pipelines for protein-ligand docking: 'docking-minimize-scoring' and 'docking-scoring-minimize'. The former approach first samples 40 conformations, perform minimization on all 40 conformations, then do scoring on the minimized samples for ranking. The latter approach first sample 40 conformations, do scoring to select the Top1 sample, then minimize it. As shown in **Supplementary Table 1**, the validity of SurfDock generated poses increase substantially after minimization using both approaches with moderate sacrifice of docking accuracy. This shows that the force field minimization stage is crucial in improving ligand validity. The performance is consistent on the unseen proteins.

While the 'docking-minimize-scoring' strategy aligns better with the need for realistic input samples for scoring, it unavoidably consumes more computational resources as it needs to optimize all samples. For practical consideration, we argue that the ‘docking-scoring-minimize’ is also useful, particularly in virtual screening scenario, for improved efficiency.

**Supplementary Table 1 | Performance of SurfDock with or without the post-docking energy minimization stage.** The best results are marked in bold, the second best are underlined

| Performance on PDBbind2020 test set | | |
|---|---|---|
| Method | %<2Å | %<2Å & PB-valid |
| SurfDock | **68.41±0.26** | 36.46±0.26 |
| SurfDock (Docking-scoring-minimize) | 62.35±0.34 | 54.45±0.26 |
| SurfDock (Docking-minimize scoring) | 68.04±0.22 | **55.00±0.13** |
| Performance on unseen proteins in PDBbind2020 test set | | |
| SurfDock | 60.88±0.33 | 30.79±0.33 |
| SurfDock (Docking-scoring-minimize) | 56.02±1.18 | 43.29±0.86 |
| SurfDock (Docking-minimize-scoring) | **62.50±0.57** | **43.75±0.56** |

### S4 Additional Results on Astex and PoseBusters Benchmark Sets

As shown in **Supplementary Fig. 2 a**, the sampling power of SurfDock on PoseBusters is consistent under different RMSD thresholds. Glide SP here is also capable of finding near-native ligand poses as SurfDock does. In **Supplementary Fig. 2 b, c**, we compare only the DL methods on Astex and PoseBusters benchmark sets. Even without the post-docking minimization stage, SurfDock has surpassed all other DL methods in both docking success rate and ligand validity. The docking-minimization-score pipeline significantly increase the ligand validity and slightly affect the overall docking success rate. This pipeline also greatly improves the generalizability of SurfDock to relatively new proteins on docking success rate and ligand validity.

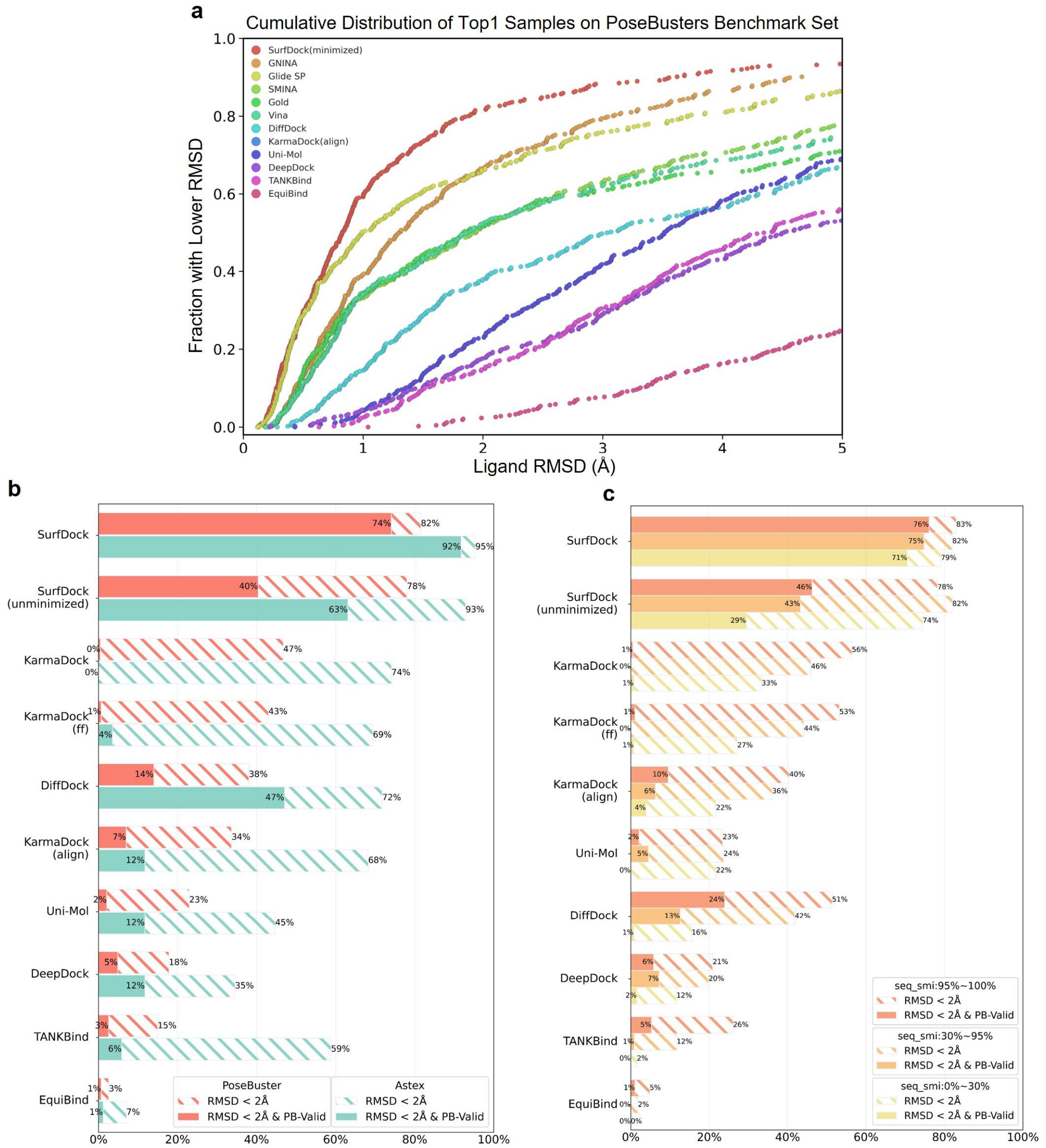


**Supplementary Fig. 2 | Comparative Performance of Different Docking Methods Across Astex and PoseBuster sets**. **a**: cumulative distribution of Top1 candidates from different methods. **b**, **c**: Comparison of different deep learning models across Astex and PoseBusters sets. We obtained the results of three versions of KarmaDock[3] by using their public model weight on https://github.com/schrojunzhang/KarmaDock.

### S5 Distribution of Rotatable Bonds and Heavy Atoms

As shown in **Supplementary Fig. 3**, the distribution of rotatable bonds and heavy atoms on PDBbind is the most challenging one, while the distribution of PoseBusters or DEKOIS2.0 is relatively simple. These distributions might help explain several experimental results:

1. The performance of SurfDock on PDBbind decreases drastically when the rotatable bonds are greater than 25, as shown in **Fig. 5**, while being consistent on PoseBusters.
2. The rescoring experiment in **Supplementary Fig. 5** shows that the sampling power of Glide SP is comparable with SurfDock, partly due to the fact that the distribution of rotatable bonds on DEKOIS2.0 is much simpler than on PDBbind, which makes the sampling task easier.

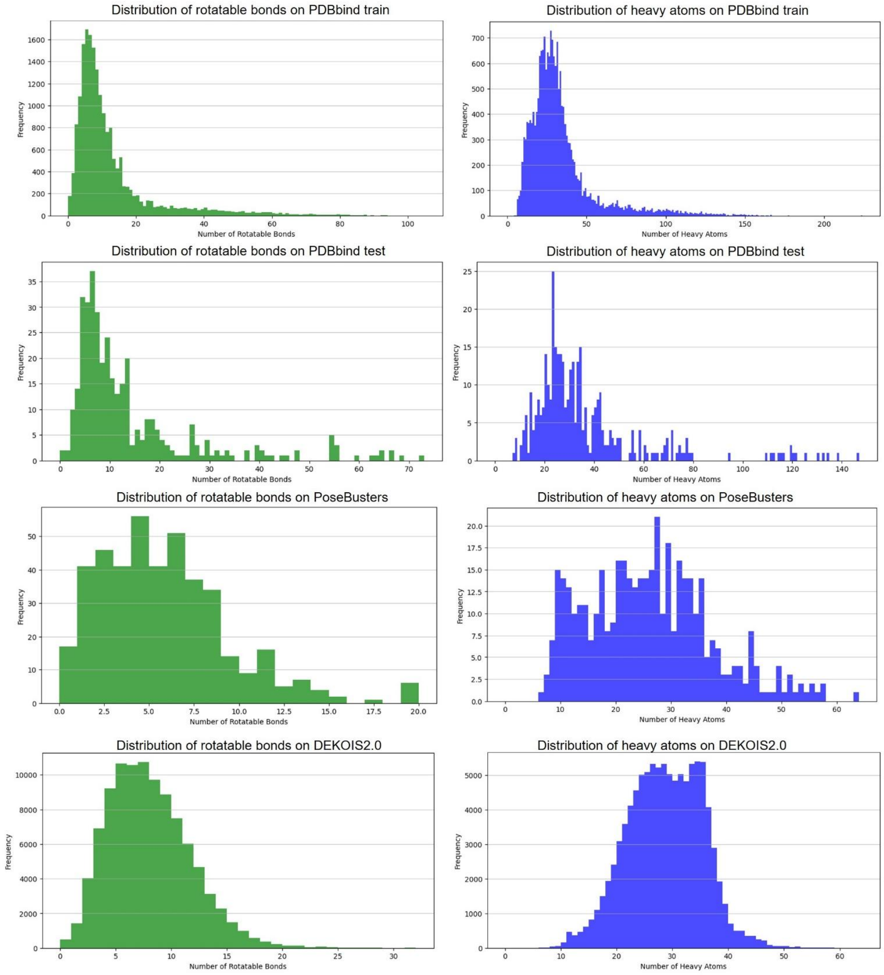


**Supplementary Fig. 3 | Distribution of the number of rotatable bonds and heavy atoms in PDBbind2020 time-split test set and PoseBusters Benchmark set**.

### S6 SurfDock Optimization Trajectories

As shown in **Supplementary Fig. 4** and **Fig.3 d**, the diffusion generative processes of selected samples with only two rotatable bonds all behave like geometry optimization processes. This shows the consistency of the diffusion generative process and the scoring module. Besides, in cases like 6i64, 6e3n, 6i66, 6npi, SurfDock can find the predicted ligand pose with confidence higher than RDKit aligned pose or even the crystal ligand pose, which is probably due to the addition of post-docking minimization stage. The incorporation of force-field features improves the rationality of generated poses, which leads to higher confident sample than even the referenced ones.

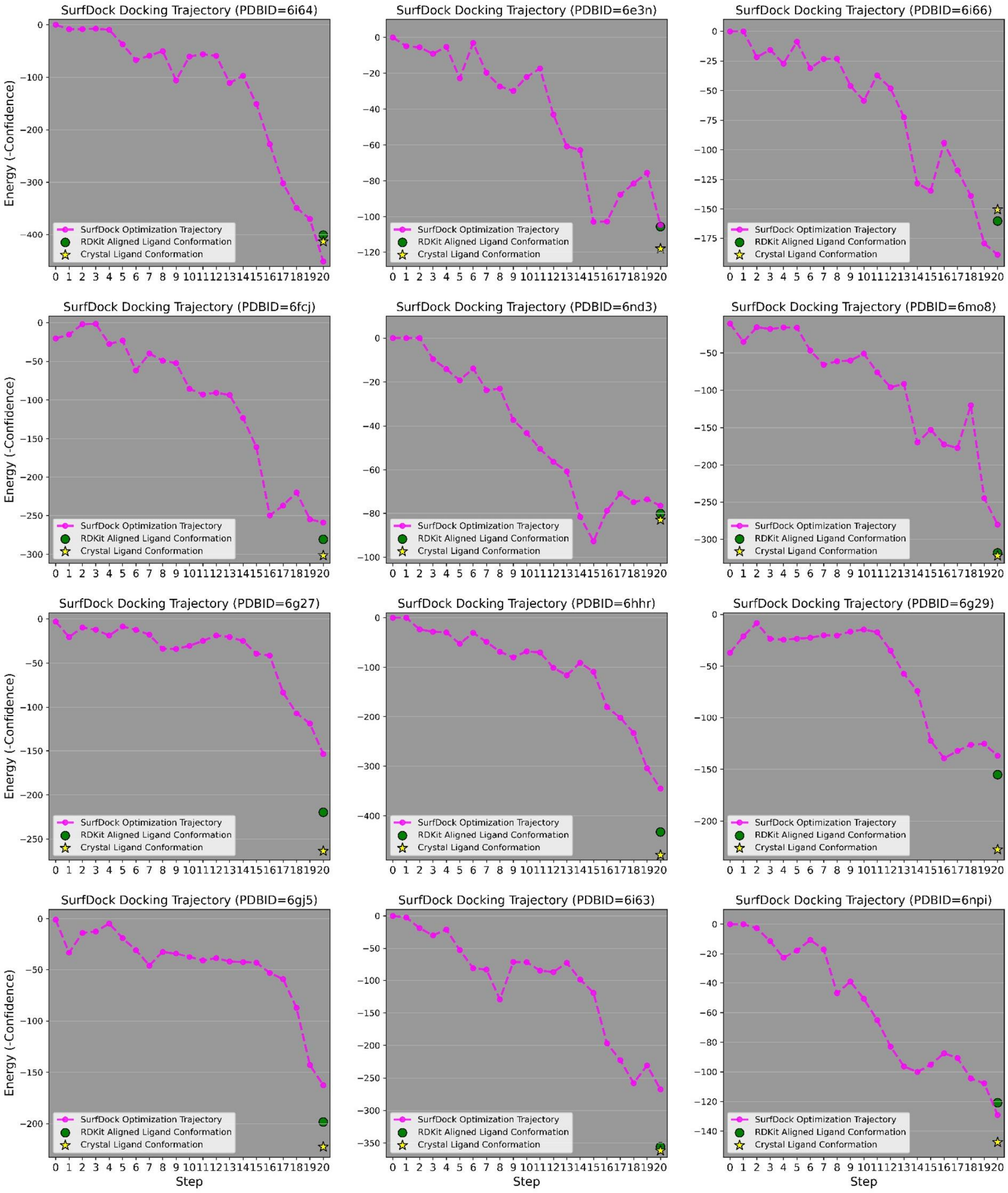


**Supplementary Fig. 4 | Examples of SurfDock Optimization Trajectories.**

### S7 Re-scoring Performance

We use the generated poses from the traditional methods and re-score them using SurfScore. It is apparent that SurfScore greatly improve the virtual screening ability of traditional docking algorithms. For comparison, a recent work RTMScore[4] achieves around 18.5 on the mean EF 1.0% metrics when combined with Glide SP, while Glide SP + SurfScore achieves around 18.7. As the focus of our research is on developing a whole pipeline considering ligand rationality, we did not further test our scoring module on more rigorous benchmarks.

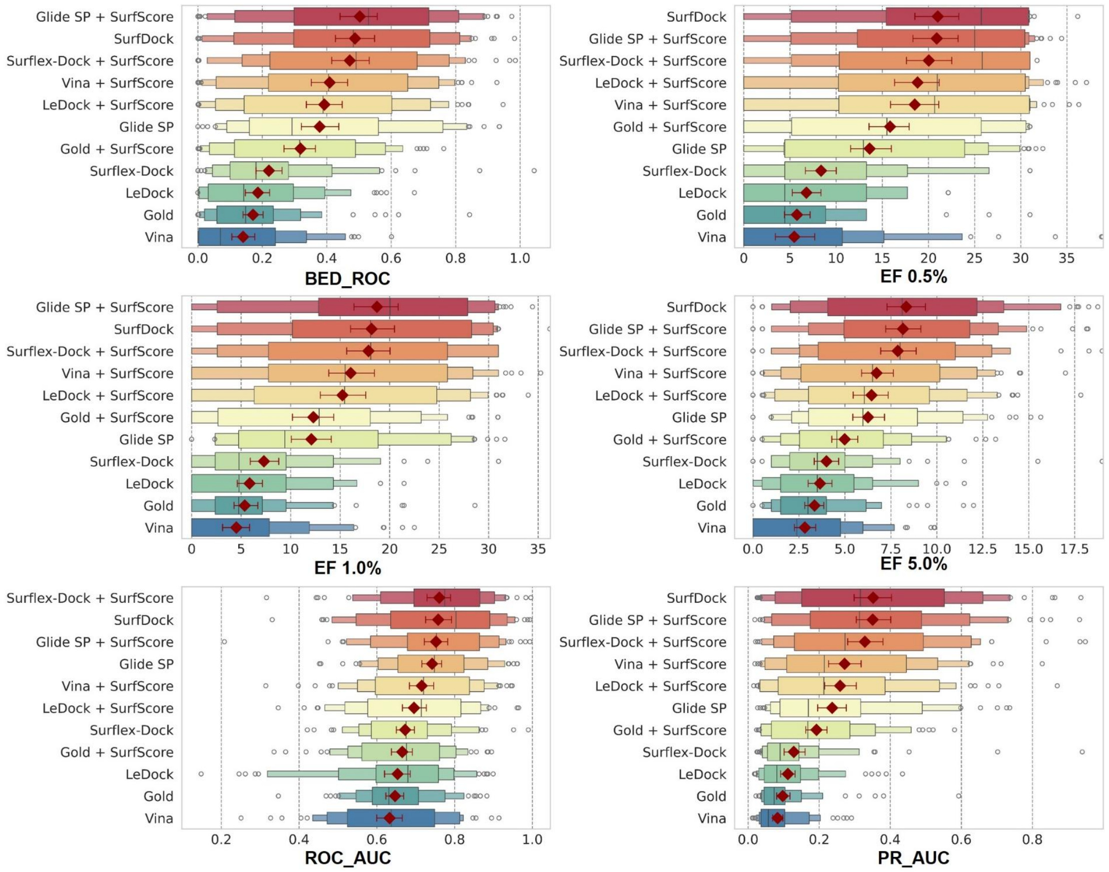


**Supplementary Fig. 5 | The Re-scoring Performance of SurfScore on DEKOIS2.0**.
